# Immunomodulatory Therapy with Glatiramer Acetate Reduces Endoplasmic Reticulum Stress and Mitochondrial Dysfunction in Experimental Autoimmune Encephalomyelitis

**DOI:** 10.1101/2021.05.04.442578

**Authors:** Tapas K. Makar, Poornachander R Guda, Sugata Ray, Sanketh Andhavarapu, Vamshi KC Nimmagadda, Volodymyr Gerzanich, Marc J Simard, Christopher T. Bever

## Abstract

Endoplasmic reticulum (ER) stress and mitochondrial dysfunction are found in lesions of multiple sclerosis (MS) and of experimental autoimmune encephalomyelitis (EAE), an animal model of MS, and may contribute to the neuronal loss that underlies permanent impairment. These pathological changes are due to the neuroinflammation that characterizes MS and the strong interplay between the ER and mitochondria. We investigated whether immunomodulatory drug glatiramer acetate (GA) can reduce these changes in the spinal cords of chronic EAE mice by using routine histology, immunostaining, and electron microscopy. EAE spinal cord tissue exhibited increased infiltration/inflammation (upregulation of proinflammatory cytokines), demyelination, mitochondrial dysfunction (increased fission, decreased fusion, and increased biogenesis), ER stress, downregulation of NAD+ dependent pathways, and increased neuronal death. GA reversed these pathological changes, suggesting that immunomodulating therapy can attenuate ER stress, mitochondrial dysfunction, apoptotic cell death, and demyelination in the CNS and increase NAD+ utilizing activities by suppressing neuroinflammation.

## 1. Introduction

Multiple sclerosis (MS) is a disease of the central nervous system (CNS), characterized pathologically by inflammation, demyelination, axonal damage and neuronal loss[1]. These events are triggered by heterogenous myelin-reactive peripheral immune cells infiltrating the blood-brain-barrier (BBB) [2]. The most widely used animal model for MS is experimental autoimmune encephalomyelitis (EAE), a murine model which presents with CNS inflammation, demyelination, axonal transection, and neurological impairment due to similar infiltration of auto-reactive cells [3–7]. The currently available disease-modifying therapies for MS have not been proven to prevent or reverse long-term neurodegeneration [8,9], although they have been shown to reduce inflammation [10]. Yet, pathological axonal damage [6] and neuronal loss [11] in the brain and spinal cord atrophy on magnetic resonance imaging (MRI) are highly correlated with long-term disability and progression more so than inflammation [12]. Therefore, MS research is currently seeking to better understand the mechanisms underlying neurodegeneration in MS in order to identify neuroprotective therapies which promote myelin repair.

Pathological studies of MS lesions have implicated mitochondrial dysfunction in the neurodegeneration in MS [13]. MRI spectroscopy shows changes in N acetyl aspartate levels consistent with decreased mitochondrial function in acute lesions [14,15]. The proximate cause of the dysfunction is relatively unknown. Reactive oxygen species (ROS) produced by mitochondria are induced in the pathogenesis of MS and EAE. Enhancement of ROS production following EAE is one of the causes of demyelination and inflammation [16,17]. As a consequence, mitochondrial ROS promotes inflammation and shifts mitochondrial dynamics towards mitochondrial breakdown [18]. Endoplasmic reticulum (ER) stress is also involved in MS lesions and in EAE animals [19,20] Upstream causes of these mitochondrial defects and ER stress are largely unknown. A possible mechanism involves the release of Ca2+ ions from the ER within the damaged neurons. This intracellular signaling mechanism ultimately induces cell death via a mitochondrial mediated mechanism [21,22]. Abnormal ER mitochondria Ca2+ transfer is observed following axotomy [23] and spinal cord injury [24]. This process not only promotes ER-mitochondria crosstalk in general by increasing the apposition of the ER and mitochondria at the mitochondria-associated membrane (MAM) [25–27], but could also promote neuronal death, thus is involved in the MS pathology [28]. Mitochondrial metabolism and ER stress are two of the most metabolically active cellular processes, playing a crucial role in regulating inflammation. Furthermore, mitochondrial dysfunction with reduced nicotinamide dinucleotide (NAD+) appears to play a role in neurodegeneration in MS pathogenesis [29–33]. Previously, we studied transgenic EAE mice with neuron specific overexpression of SIRT1, an NAD dependent enzyme, and found evidence supporting a protective role of SIRT1 in EAE [34] prevented or altered the phenotype of inflammation in spinal cords; as a result, demyelination and axonal injury were reduced. SIRT1 also regulates mitochondrial function [35], ER stress [36] and inflammation [37]. This has shifted research focus towards the effects of inflammation on mitochondrial dysfunction, ER stress and NAD associated SIRT1 pathway.

Glatiramer Acetate (GA), commonly known as copaxone, is an approved immunomodulatory disease-modifying drug for MS treatment [38]. GA mediates remyelination by altering T-cell differentiation and inducing a shift towards Th2/3 cells and neurotrophic factors that can penetrate the blood-brain-barrier (BBB) and accumulate in the CNS [39–42]. To date, the effects of GA on ER stress, mitochondrial dysfunction and NAD+ SIRT1 in the CNS of EAE mice has not been studied, so we set out to investigate this. Using routine histology, immunohistochemistry, western blotting, and electron microscopy, we discovered that GA can regulate ER stress, mitochondrial function and dynamics, and NAD+ dependent pathways by inhibiting neuroinflammation. Our findings elucidate that targeting neuroinflammation associated with ER stress, mitochondrial dysfunction and NAD regulated mechanism through a peripherally acting drug can be neuroprotective in MS.

## 2. Methods

### 2.1 Animals and Drug

Female C57Bl/6J mice were obtained from The Jackson Laboratory (Bar Harbor, ME, USA). Mice were housed in our facilities under pathogen-free conditions at the University of Maryland, School of Medicine, Baltimore. Mice were maintained and treated following NIH guidelines and the Institutional Animal Care and Use Committee-approved protocol. EAE was induced in eight week old female C57Bl/6 mice with 0.2 mg of the myelin oligodendrocyte glycoprotein (MOG) 35-55 peptide in complete Freund’s adjuvant (CFA) followed by pertussis toxin injections as we have described previously [34]. Mice were scored daily by blinded raters using a standard impairment scale as described previously[34]. GA (TEVA Neuroscience) was administered subcutaneously at a dose of 125µg/mouse/day in a 200µl vehicle of phosphate-buffered saline (PBS). The mice were randomized to vehicle-treated EAE (EAE), GA-treated EAE (EAE+GA) and vehicle-treated age matched normal female mice. All mice received a single injection daily starting from the day of disease onset among the EAE mice (Score ≥ 1).

### 2.2 Tissue Pathology

Mice were euthanized and spinal cords and brains were removed and prepared for histology as described previously [34]. Immunohistochemical and immunofluorescence studies were carried out as described previously [34]. The primary antibodies used are listed in Table 1.

**Table 1:**
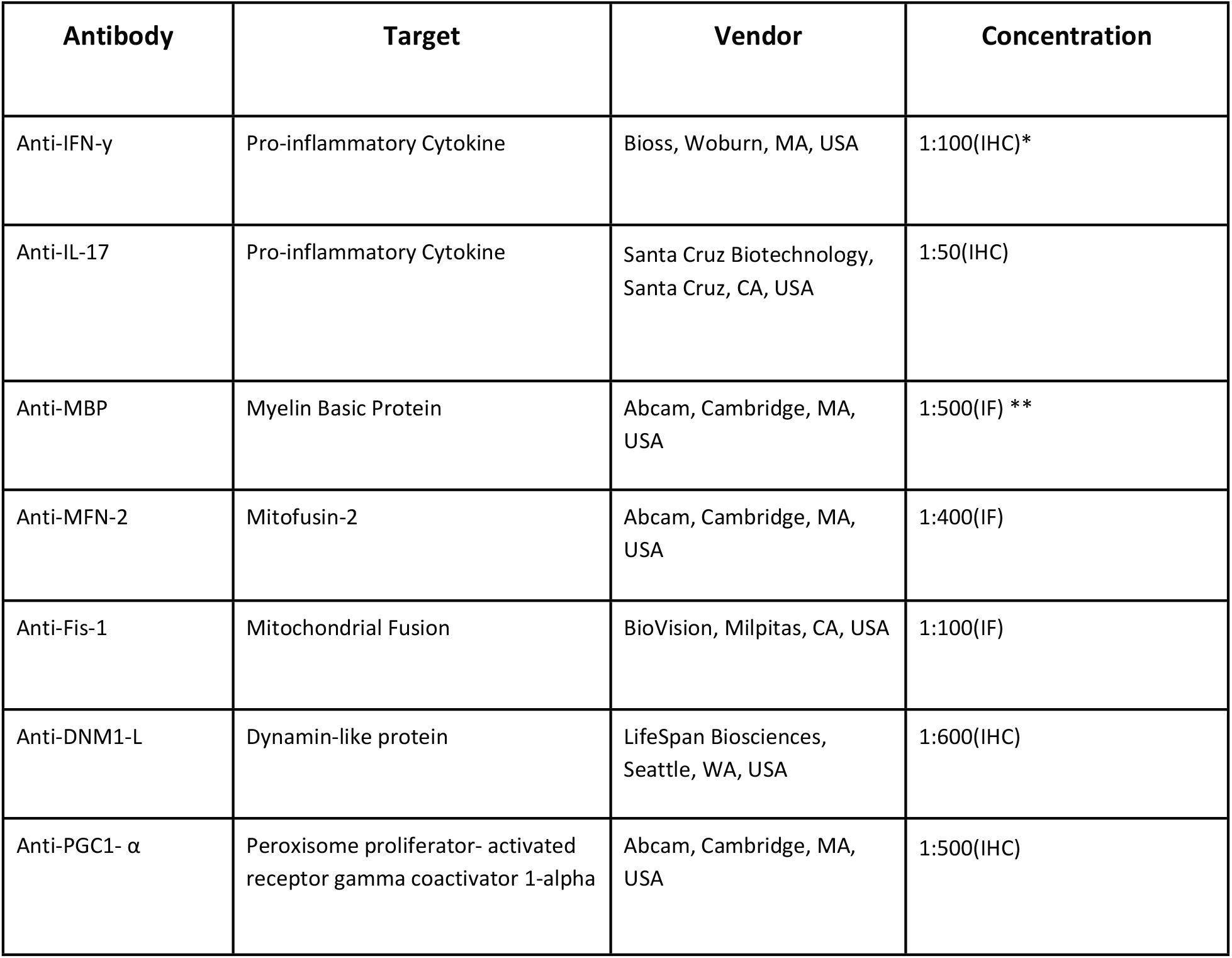
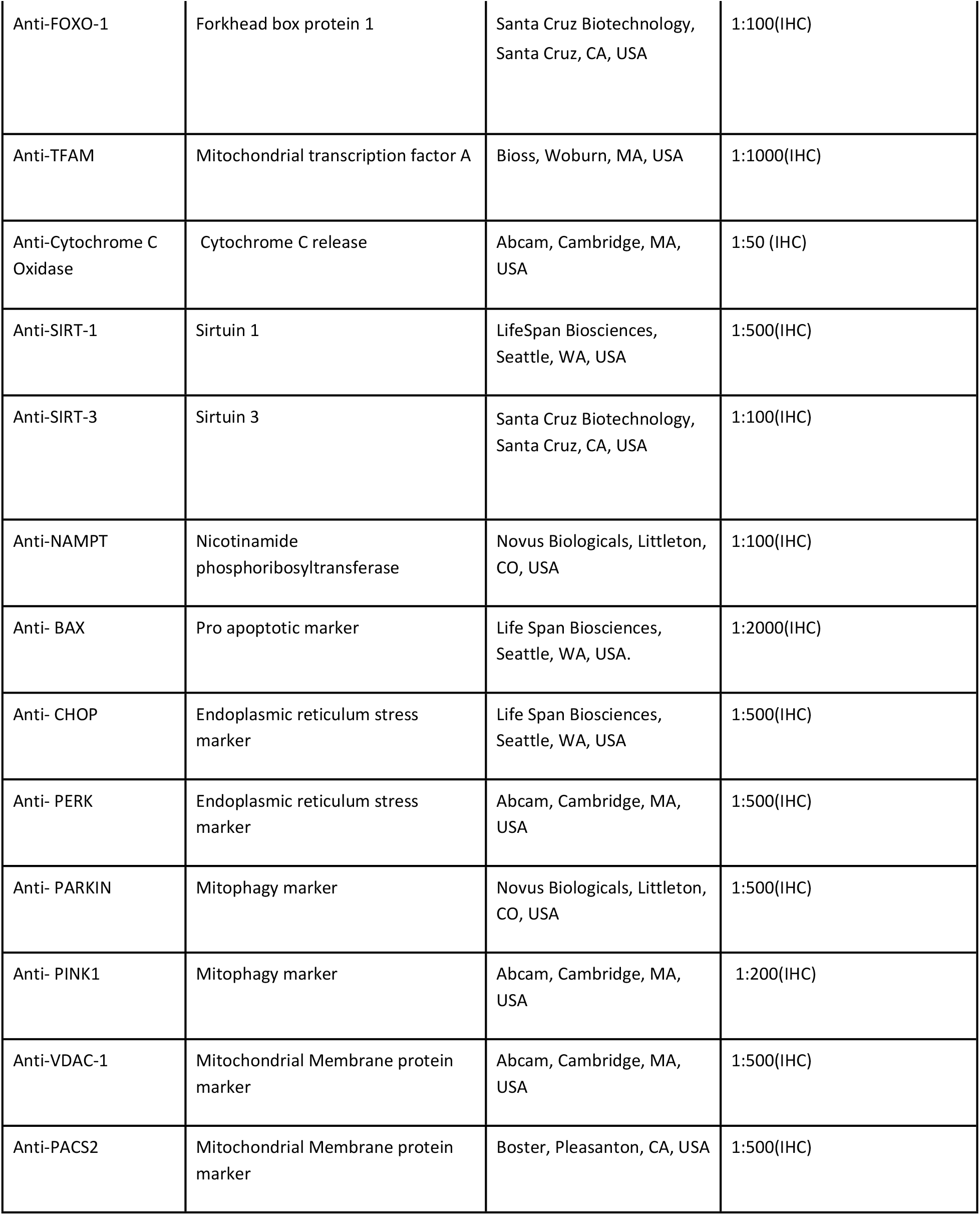
List of antibodies used in the study.

| Antibody | Target | Vendor | Concentration |
| --- | --- | --- | --- |
| Anti-IFN-γ | Pro-inflammatory Cytokine | Bioss, Woburn, MA, USA | 1:100(IHC)* |
| Anti-IL-17 | Pro-inflammatory Cytokine | Santa Cruz Biotechnology, Santa Cruz, CA, USA | 1:50(IHC) |
| Anti-MBP | Myelin Basic Protein | Abcam, Cambridge, MA, USA | 1:500(IF) ** |
| Anti-MFN-2 | Mitofusin-2 | Abcam, Cambridge, MA, USA | 1:400(IF) |
| Anti-Fis-1 | Mitochondrial Fusion | BioVision, Milpitas, CA, USA | 1:100(IF) |
| Anti-DNM1-L | Dynamin-like protein | LifeSpan Biosciences, Seattle, WA, USA | 1:600(IHC) |
| Anti-PGC1- $\alpha$ | Peroxisome proliferator- activated receptor gamma coactivator 1-alpha | Abcam, Cambridge, MA, USA | 1:500(IHC) |
| Anti-FOXO-1 | Forkhead box protein 1 | Santa Cruz Biotechnology, Santa Cruz, CA, USA | 1:100(IHC) |
| Anti-TFAM | Mitochondrial transcription factor A | Bioss, Woburn, MA, USA | 1:1000(IHC) |
| Anti-Cytochrome C Oxidase | Cytochrome C release | Abcam, Cambridge, MA, USA | 1:50 (IHC) |
| Anti-SIRT-1 | Sirtuin 1 | LifeSpan Biosciences, Seattle, WA, USA | 1:500(IHC) |
| Anti-SIRT-3 | Sirtuin 3 | Santa Cruz Biotechnology, Santa Cruz, CA, USA | 1:100(IHC) |
| Anti-NAMPT | Nicotinamide phosphoribosyltransferase | Novus Biologicals, Littleton, CO, USA | 1:100(IHC) |
| Anti- BAX | Pro apoptotic marker | Life Span Biosciences, Seattle, WA, USA. | 1:2000(IHC) |
| Anti- CHOP | Endoplasmic reticulum stress marker | Life Span Biosciences, Seattle, WA, USA | 1:500(IHC) |
| Anti- PERK | Endoplasmic reticulum stress marker | Abcam, Cambridge, MA, USA | 1:500(IHC) |
| Anti- PARKIN | Mitophagy marker | Novus Biologicals, Littleton, CO, USA | 1:500(IHC) |
| Anti- PINK1 | Mitophagy marker | Abcam, Cambridge, MA, USA | 1:200(IHC) |
| Anti-VDAC-1 | Mitochondrial Membrane protein marker | Abcam, Cambridge, MA, USA | 1:500(IHC) |
| Anti-PACS2 | Mitochondrial Membrane protein marker | Boster, Pleasanton, CA, USA | 1:500(IHC) |

### 2.3 TUNEL assay for apoptotic cell death

ApopTag Peroxidase Kit (Chemicon International, Temecula, CA) was used to assess the extent of cell death in the spinal cord sections of EAE animals and GA treated animals. Briefly, all the slides were deparaffinized with xylene and rehydrated through graded alcohols to water, washed in 0.01 M PBS then pretreated with a protein-digesting enzyme for 15 min and then washed with water for 2 min. slides were treated with 3% (v/v) hydrogen peroxide for 5 min followed by washing with PBS. Terminal deoxynucleotidyl transferase (TdT) enzyme was added to the pre-equilibrated spinal cord sections and incubated for 1 h at 37°C. Stop-buffer was added to the slide and agitated for 15 sec followed by 10 min incubation at room temperature. After washing three times with PBS for 1 min each, anti-digoxigenin peroxidase conjugate was added to the slides and incubated for 30 min. After slides were washed twice with PBS, freshly prepared peroxidase substrate 3,3’-diaminobenzidine was added to the slides and kept for 6 min and then slides were washed with water three times. Slides were counterstained with 0.5% (w/v) DAB for 5 min followed by washing with water and then 100% n-butanol. After 10 min, cells were dehydrated in xylene for 2 min and then mounted with glass coverslip. Experiments were conducted in triplicates and the ApopTag-positive cells was determined by counting cells under light microscopy

### 2.4 Analysis of histological images using Image J

Image J was used for histological quantification by a blinded observer. Cell infiltration was quantitated by counting the number of positive quadrants with inflammation, and then expressed as a percentage over the total number of quadrants examined in the histogram as reported previously[34]. Demyelination was quantitated using LFB staining and MBP staining as described previously[34]. The cell labeling experiments (IL-17, IFN-γ, Cytochrome C, PINK-1, PERKIN, DNM1-L, PGC-1alpha, TFAM, CHOP, PERK, PACS-2, VDAC-1, NAMPT, SIRT-1, SIRT-3, FOXO-1, BAX, MBP, NFM-2, Fis-1) were quantified based on the number of positive cells/field (200X or 400X). All fields covering the entire white matter (10-12 fields/section) and gray matter (5-7 fields/section) were analyzed from each spinal cord. The cell counting and data analysis were performed by an examiner blinded to treatment assignment.

### 2.5 Electron Microscopy

Spinal cords were excised and fixed with a solution of 2% paraformaldehyde, 2.5% glutaraldehyde, in 0.1M PIPES buffer (pH 7) at 4°C overnight. Specimens were washed with 0.1M PIPES buffer (pH 7), treated with 50mM glycine in 0.1 M PIPES buffer for 15 minutes and washed again with 0.1M PIPES buffer. Tissue pieces were then post-fixed in 1% osmium tetroxide, 1.5% potassium ferrocyanide in 0.1 M PIPES buffer for 60 min, washed and followed by en bloc staining with 1% (w/v) uranyl acetate for 60 min. After washing, specimens were dehydrated using a serial graded ethanol solution (30%, 50%, 70%, 90% and100%) and then 100% acetone. After dehydration, specimens were infiltrated and embedded in Araldite resin (Electron Microscopy Sciences, PA) following manufacturer’s recommendation. Ultra-thin sections∼70nm thickness were cut on a Leica UC6 ultramicrotome (Leica Microsystems, Inc., Bannockburn, IL) and collected onto copper grids and examined in an FEI Tecnai T12 transmission electron microscope (FEI. Co., Hillsboro, OR) operated at 80kV. Digital images were acquired by using a bottom mount CCD camera (Advanced Microscopy Techniques, Corp, Woburn, MA) and AMT600 software. A minimum of 3 different grids were examined for each animal from each group (N=5). All the grids were examined at x1100 for identification of white matter and gray matter and then examined at higher magnification for mitochondrial structure and endoplasmic reticulum integrity (x6,500 and x 11,000).

### 2.6 Statistical analysis

Data analysis was performed using Prism software (Graph Pad, San Diego, CA) and groups were compared using one-way analysis of variance (ANOVA) with Fisher’s protected least significant difference (PLSD) post hoc test at a 95% confidence interval. All results were presented as mean ± standard error of mean (SEM) of separate experiments (n ≥ 5). Differences were considered significant at *p* ≤ 0.05.

### 2.7 Data availability

The primary data upon which this manuscript is based is available from the corresponding author upon reasonable request. This study was not pre-registered.

## 3. Results

### 3.1 GA treatment improves clinical score, reduces demyelination, and suppresses inflammation in EAE mice

<u>Clinical Score:</u> All mice developed clinical symptoms at about post immunization day (p.i.d.) 9 (± 2.0) and without treatment average clinical scores remained elevated through p.i.d. 30 (Figure 1A1). GA treatment started at the time of symptom onset reduced the severity of disease within three days and scores remained significantly lower than the untreated EAE group with average clinical scores over days 10 to 30 of 0.57± 0.06 compared with 1.8 ± 0.15; P < 0.0001 (Figure 1A2).

<u>Demyelination</u> was assessed in sections of spinal cord using Luxol Fast Blue (LFB) staining (Figures 1B1-1B3). Quantitative analysis of the extent of demyelination shows 40.99 ± 2.7 % of white matter with myelin loss (demyelination) in EAE as compared to 8.62 ± 2.4 % in EAE+GA (P < 0.0001) (Figure 1B4). These results were confirmed using immunohistochemical analysis with antibodies against myelin basic protein (MBP) (Figures 1B5-1B7). Quantitative analysis of the percentage of white matter with MBP loss showed significant loss in EAE that was reduced by GA treatment (Figure 1B8). The changes in demyelination were confirmed qualitatively by transmission electron microscopy at 6500x (Figure 1B9-1B11).

<u>Production of pro-inflammatory cytokines</u> was examined by immunohistochemical staining for interleukin-17 (IL-17) and interferon-gamma (IFN-g) (Figure 1C). The number of cells expressing IL-17 was increased in EAE P>0.05 but reduced by GA treatment, P<0.05 (Figure 1C4). The number of cells expressing IFN-g was increased in EAE P<0.001 but reduced by GA treatment P<0.001 (Figure 1C8).

<u>Inflammatory cell infiltration</u> was examined in H&E stained sections of spinal cord from mice euthanized on day 30 (Figure 1C9-C11). While normal mice did not have cellular infiltration, EAE mice had 47.09 ± 1.86 % of white matter with inflammation compared to 12.95 ± 2.34 % in EAE+GA (Figure 1C12, P<0.0001).

**Fig 1: Glatiramer Acetate treatment improves clinical score, reduces demyelination, and suppresses inflammation in EAE mice**. (A) The clinical severity of EAE disease was reduced by GA treatment. (1A) Graph of mean EAE scores for both groups of mice over a 30-day period. (1B) Mean score for all animals in each group during the disease phase (days 10-30). The mean score was significantly lower in the GA treated EAE (EAE+GA) mice (P<0.0001) compared to untreated EAE mice. N=20/group, t-test. (B) Demyelination in EAE was reduced by GA treatment. Demyelination was assessed on LFB-stained (B1 – B3) and MBP stained (B5 – B7) sections of spinal cord white matter. Original magnification was x400. B4 and B8 are graphs of the number of quadrants with demyelination expressed as a percentage of the total number of quadrants examined (n=4), with statistics based on the t-test. B9 – B11 show transmission electron microscographs (TEM) at 6500x magnification in normal mouse (B9) untreated EAE mice (B10) and EAE with GA treatment (B11). These images are representative of at least 3 grids from each of five mice in each group. (C) Inflammation in EAE was reduced by GA treatment. Inflammation was assessed in spinal cord white matter by (C1 – C4) cells staining positively for antibodies to IL-17 and (C5 – C8) cells staining positively for IFNγ. Original magnification was x400. Hematoxylin and eosin (H&E) stained sections show infiltration of mononuclear cells in white matter of EAE (C10) and EAE+GA (C11) mice. However, the number of inflammatory pockets and inflammatory cells is fewer in EAE+GA compared to EAE. Normal (C9) mice show no inflammatory infiltrates. (C12). The number of positive quadrants with inflammation was scored and expressed as a percentage of the total number of quadrants (H&E; n=4). Bar graphs show cell counts and statistical comparisons based on one-way ANOVA.

### 3.2 GA reduces ER Stress in EAE spinal cord

We examined ER stress by assessing cells expressing CCAAT-enhancer-binding protein homologous protein (CHOP), which is induced by ER stress and is a mediator of apoptosis [43]. The number of cells positive for CHOP was increased in EAE but decreased in EAE mice treated with GA (Figure 2A1-2A3). Figure 2A4 shows the quantitative analysis of CHOP expression, which was significantly decreased in EAE+GA compared to EAE (P<0.01). Next we examined ER stress in sections of the spinal cord using antibodies to protein kinase RNA-like endoplasmic reticulum (PERK) a component of mitochondria-associated ER membranes[44]. The number of cells positive for the PERK was increased in EAE but decreased in EAE mice treated with GA (Figure 2B1 - 2B3). Figure 2B4 shows the quantitative analysis of GFAP expression, which was significantly decreased in EAE+GA compared to EAE. Finally, we examined the structural integrity of the endoplasmic reticulum using transmission electron microscopy with 6500x magnification and found ER disruption in EAE that was not present in normal or GA treated EAE (Figure 2C1-2C3).

**Fig 2: GA reduces ER stress in EAE spinal cord**. Endoplasmic reticulum stress was assessed with antibodies to CHOP (A1 – A4) and PERK (B1 – B4). Panels 1 – 3 show representative staining patterns in each treatment group for each antibody. Original magnification: X400. Panel 4 in each case shows a graph of the mean number of positive cells per field for each treatment group (CHOP : n = 4; PERK: n = 4) with intergroup comparisons based on a one-way ANOVA. Transmission electron microscopic examination (C1 – C3) showed vesiculated endoplasmic reticulum, irregularly arranged and disrupted endoplasmic reticulum (arrows) in untreated EAE (C2) while regular and parallel organized endoplasmic reticulum in normal (C1) and GA treated EAE mice (C3).

### 3.3 GA treatment improves mitochondrial function, fission/fusion, and biogenesis in EAE spinal cord

<u>Mitochondrial integrity</u> was assessed by immunohistochemical and immunofluorescent staining. Function was assessed using expressions of PINK-1 and PARKIN. PINK-1 accumulates on poorly functioning (depolarized) mitochondria allowing PARKIN to bind, targeting the mitochondria for autophagy [45]. PINK-1 and PARKIN were increased in untreated EAE but not in EAE treated with GA (Figure 3A1-3A3 and 3B1-3B3), confirming lost mitochondrial integrity in EAE and revealing that GA can restore this. Figures 3A4 and 3B4 shows the quantitative analysis of PGC1-α expression

<u>Mitochondrial biogenesis</u> was examined in spinal cords for cells expressing PGC1-α, a mediator of mitochondrial biogenesis[46], by immunohistochemistry (Figure 3C1 - 3C3). Figure 3C4 shows the quantitative analysis of PGC1-α expression, which was significantly reduced in EAE mice P<0.01, but restored back to normal levels in EAE mice treated with GA (P<0.01).

<u>Morphological changes</u> in mitochondria were assessed directly by transmission electron microscopy at 6500X magnification (Figure D1-D3). Mitochondrial fission, cristae damage and loss, mitochondrial membrane damage and disappearance, and changes in size and shape were seen in EAE mice but not in the tissues from normal mice or EAE mice treated with GA. Mitochondrial fusion was not seen.

<u>Mitochondrial fusion</u>, which is reduced under conditions of ER stress, was assessed by immunohistofluorescence staining (Figure 3E1-3E3) and western blot analysis for MFN-2, a marker of fusion[47]. The number of cells positive for the MFN-2 was reduced in EAE (P<0.05) but not in EAE mice treated with GA compared to normal mice (P<0.05) (Figure 3E4).

<u>Mitochondrial fission,</u> which is increased under conditions of mitochondrial stress, was assessed by immunofluorescence staining for FIS-1[48] and both immunohistochemistry and western blot analysis for DNM1-L (Figure 3F1-3F3 and 3G1-3G3, 3G5)[49]. The number of cells expressing FIS-1 did not change in the EAE mice but significantly decreased in EAE mice treated with GA compared to control EAE mice (P<0.05) (Figure 3F4). The number of cells expressing DNM1-L significantly increased in the EAE mice (P<0.01) but decreased to normal levels in EAE+GA (P<0.01) (Figure 3G4).

**Fig 3: GA treatment improves mitochondrial function, fission/fusion, and biogenesis in EAE mice**. (A,B) EAE caused loss of mitochondrial integrity that was reduced by GA treatment. Mitochondrial integrity was assessed by staining for PINK-1 (A1 – A4) and staining for PARKIN (B1 – B4). Panels 1 – 3 show representative staining patterns in each treatment group for each antibody. Original magnification: X400. Panel 4 in each case shows a graph of the mean number of positive cells per field for each treatment group (n = 4) with intergroup comparisons based on a one-way ANOVA. (C,D) Changes in mitochondrial biogenesis and morphology in EAE are reduced by GA treatment. Mitochondrial biogenesis was assessed with antibody to PGC1-α, a regulator of mitochondrial biogenesis (C1 – C4) Panels 1 – 3 show representative staining patterns in each treatment group for each antibody. Original magnification: X400. Panel 4 shows a graph of the mean number of positive cells per field for each treatment group (n = 3; n = 4) with intergroup comparisons based on a one-way ANOVA. Mitochondrial morphology was assessed by transmission electron microscopy (TEM) at 6500x magnification. Morphological changes were observed in mitochondria in EAE (D2) that were reduced by treatment (D3) compared to the mitochondria in normal mice (D1). In EAE many of the mitochondria showed mitochondrial membrane disruption with disruption or loss of cristae and changes in mitochondrial size and shape with increased fission. The results shown are typical of at least 3 grids for each mouse (n = 5). (E-G) EAE caused changes in mitochondrial dynamics that were reduced by GA treatment. Mitochondrial fusion was assessed by staining with antibody to Mitofusin – 2 (MFN-2) (E1 – E4) and mitochondrial fission was assessed with antibody to FIS-1 (F1 – F4) and DNM1-L (G1 – G4). Panels 1 to 3 show representative staining patters in each treatment group with each antibody. Original magnification was x400. Panel 4 in each case shows a graph of the mean number of positive cells per field for each treatment group (MFN-2, n = 4; FIS-1, n = 3; DNM1-L, n = 3) with intergroup comparisons based on a one-way ANOVA. (E5) Western Blot analysis of mitochondrial fraction of spinal cord shows an increase in MFN-2 in the EAE+GA mice compared to EAE and Normal. (G5) Western Blot analysis of mitochondrial fraction of spinal cord shows a decrease in DNM1-L in the EAE+GA mice compared to EAE and Normal.

### 3.4 GA regulates changes in the mitochondria associated membrane (MAM) and increases activity of the NAD+ dependent pathway in EAE spinal cord

Mitochondria and the endoplasmic reticulum interact through a specialized domain on the ER called the mitochondria-associated membranes (MAM). Several proteins are located in MAMs, including PACS-2, VDAC-1[50,51]. We examined PACS-2 and VDAC-1 expressions in EAE using immunohistochemistry and evaluated the effect of GA treatment (Figure 4A1-4A3 and 4B1-4B3). Numbers of cells expressing PACS-2 and VDAC-1 (Figure 4A4 and 4B4) were increased in EAE (P<0.05) but returned toward normal in GA treated mice (P<0.05).

**Fig 4: GA regulates changes in mitochondria associated membrane (MAM) associated with EAE in the spinal cord**. Antibodies to PACS2 (A1–A4) and VDAC-1 (B1–B4) were studied as markers of the MAM. (A1 – A3 and B1 – B3) show the antibody staining in each treatment group with each antibody at an original magnification of x400. (A4 and B4) show quantitation of staining in each treatment group with comparisons by one-way ANOVA (n=5).

Because of the importance of NAD regulation in mitochondrial function and cell survival[52] we examined expression of Nicotinamide phosphoribosyltransferase (NAMPT), the rate limiting enzyme in NAD biosynthesis[53], and Sirtuins 1 and 3 which are NAD dependent protein deacetylases[54] which play a role in mitochondrial dynamics. Staining for NAMPT, SIRT-1 and SIRT-3 and was reduced in EAE compared to normal (Figures 5A2, 5B2, 5C2). Treatment with GA partially restored levels of NAMPT, SIRT-1 and SIRT-3 (Figures 5A3, 5B3, 5C3). Figures 5A4, 5B4 and 5C4 show the quantitative analysis of NAMPT, SIRT-1 and SIRT-3 expressions.

**Fig 5: GA increased the activity of NAD-dependent pathways**. NAD+ dependent pathways were assessed with antibodies to NAMPT (A1 – A4), Sirt-1 (B1 – B4), and Sirt-3 (C1 – C4). For each antibody, panels 1 – 3 show typical staining in each treatment group at an original magnification of x400 and panel 4 is a graph of the quantification of positive cells in the spinal cord with comparisons made using one-way ANOVA (n = 4).

### 3.5 GA reduces apoptosis in EAE spinal cord

Because of mitochondrial involvement in programmed cell death we assessed apoptosis in EAE using the TUNEL assay. TUNEL cells were significantly increased in EAE (P<0.001) but lower in EAE+GA spinal cords (P<0.001) (Figure 6A1-6A3). Figure 6A4 shows the quantitative analysis of TUNEL positive cells in the gray matter of these spinal cords. To evaluate activation of the apoptosis pathway we examined the expression of FOXO-1, a transcription factor regulated by AKT and implicated in apoptosis [55] in part by the regulation of Bax, an activator of apoptosis[56]. FOXO-1 expressing cells were increased in EAE (P<0.05) but normal in GA treated mice (P<0.001) (Figure 6B1-B4). BCL-2 expressing cells were increased in EAE (P<0.05) but lower after GA treatment (P<0.05) (Figure 6C1-4). Staining for cytochrome-C expression in spinal cord sections was used as a measure of apoptosis triggered by decreased mitochondrial integrity[57](Figure 6A1-6A3). Cytochrome C expressing cells were increased in EAE (P<0.001) but was reduced back to normal levels in GA treated mice (p<0.001) (Figure 6A4).

**Fig 6: GA reduces apoptosis in EAE spinal cord**. Apoptosis was assessed in spinal cord tissue with in situ TUNEL staining (A1 – A4) and with antibodies to FOXO-1 (B1 – B4), BAX (C1 – C4), and Cytochrome C (D1-D4). Panels 1 – 3 show typical staining patterns for each treatment group at an original magnification of x 400 while panels 4 provide quantitation of numbers of positive cells in each group (n = 4) with comparisons using one-way ANOVA.

## 4. Discussion

In this study, we investigated for the first time the effects of GA, a polypeptide-based approved drug for the treatment of MS, on ER stress and mitochondrial dysfunction and NAD related pathways, along with neuroinflammation and demyelination, in the spinal cord of EAE mice. Our principle findings are: (i) as expected, GA improved clinical score, reduced inflammatory activity, and promoted remyelination; (ii) GA reduced ER stress; (iii) GA improves mitochondrial function, reduces mitochondrial fission, increases mitochondrial fission, and increases mitochondrial biogenesis; (iv) GA regulated changes in mitochondrial associated membranes; (v) GA increases activity of the NAD+ dependent pathway; and (vi) GA reduces apoptosis.

Research thus far has demonstrated that GA exerts its immunomodulatory effects by altering T-cell differentiation through promotion of Th2-polarized GA-reactive CD4+ T-cells [58]. Furthermore, induction of Th2 cells in the periphery during GA treatment leads to reduced inflammation, and in turn promotes remyelination and neuronal survival [59]. Figure 1 of our research confirms these findings. Specifically, we showed that GA downregulated expression of IFN-y, which promotes myelin damage by stimulating inflammation. It is important to note that GA is degraded in the periphery and cannot cross the blood brain barrier, so the spinal cord findings that we present demonstrate GA’s in situ bystander effect [42]. Furthermore, GA treatment increased Th2 cytokine IL-10 expression. It is recently reported that GA treatment causes a switch from pro-inflammatory microglia and astrocytes to anti-inflammatory phenotypes of those CNS cells that are involved in MS pathogenesis [60].

ER stress is involved in various intracellular physiological functions such as protein folding, calcium homeostasis, lipid metabolism, cell differentiation, and protein translocation [61]. ER stress is characterized by the accumulation of misfolded proteins, resulting in chronic perturbations to ER homeostasis [62]. Increasing evidence suggests that ER stress, which is caused by the accumulation of unfolded or misfolded proteins in the ER, plays a role in inflammatory diseases, including MS and EAE [63].

The unfolded protein response (UPR) is an evolutionary conserved process that is activated in order to restore ER homeostasis by correcting protein-folding machinery. The UPR has three main arms led by ER-transmembrane proteins: PERK, IRE1, and ATF6. In this study, we focused specifically on PERK and its downstream apoptotic gene CHOP. Although ER stress initially acts as self-preservation, chronic ER stress and activation of the UPR leads to cellular apoptosis [64]. It is widely known that the UPR is activated in both MS and EAE lesions, induced by elevated levels of proinflammatory mediators, and contributes to disease progression [65]. Specifically, previous studies have demonstrated that EAE, including spinal cord tissue, exhibits upregulated levels of p-PERK and CHOP in oligodendrocytes, T cells, astrocytes, and macrophages/ microglia [50,65–67]. We confirmed the increases in PERK and CHOP using immunostaining and showed that they were reversed by immunomodulating treatment with GA. In our study, we found that GA reversed the activation of PERK and CHOP in the EAE model, and this was supported by our electron microscopy that presented restored ER structure similar to that of wild type mice. We suggest that this is due to GA’s ability to mediate neuroinflammation, specifically IFN-y. IFN-y has previously been shown to induce PERK activation and its downstream translation initiation factor 2 (eIF2α), and IFN-y induced apoptosis in rat oligodendrocytes is associated with ER stress [68]. Targeting the PERK-eIF2α pathway has been reported as an ideal strategy for protecting oligodendrocyte protection in MS, and we show for the first time that GA may be able to [69]. Other chemical compounds that have been shown to activate this pathway and are neuroprotective in EAE and MS include salburnal [70] and guanabenz [71]. Future studies with GA should investigate its cytoprotective effects on other branches of the UPR in specifically oligodendrocytes and neurons. It is demonstrated that ER stress generated in murine astrocytes encourages PERK-dependent inflammatory signaling in vitro, suggesting that astrocytes themselves are potential contributors to neurotoxic inflammation in the face of ER dysfunction [66,72]. Particularly we provided evidence of the link between the effector cytokine molecules in regulating ER stress by GA treatment in EAE mice. Furthermore, these studies suggest that controlling balance of ER stress and inflammatory response can serve as an important therapeutic mechanism of GA for regulating EAE and perhaps MS.

Mitochondrial dysfunction is a pathological hallmark in EAE and MS lesions, and it is well associated with ER stress [19,20]. Therefore, we were interested in examining changes in mitochondrial dynamics to determine if GA would reverse such changes, possibly through attenuation of the ER-stress induced UPR. The PERK-ATF4-CHOP pathway exerts regulatory effects on the expression of Parkin, a critical regulator in mitochondrial dynamics (Sarrabeth Stone 2015). Parkin plays a role in mitochondrial dynamics [73,74], bioenergetics [75,76], and mitophagy [45]. Parkin also is known to modulate MAMs, which we assessed by staining for PACS2 and VDAC1, to maintain calcium transfer between the ER and mitochondria [75]. Haile et al recently showed that during MS progression, ER stress is strongly associated with the upregulation of Rab32, a GTPase that regulates MAMs, and contributes to neuronal death. We found that both the PINK1/Parkin pathway and MAMs was upregulated in the EAE model as found previously [77], and GA reversed this, likely by suppressing the PERK branch of the UPR.

The UPR, PINK1/Parkin pathway, and MAMs all play a role in mitochondrial fission/fusion processes. ER-stress induced PERK regulates the Drp1–Fis1 complex through control of the adaptor protein AKAP121. PINK1 and Parkin promote mitochondrial fission via a Drp1 mediated mechanism [78]. Parkin also negatively regulates mitochondrial fusion via MFN2 by ubiquitination [79] and interestingly, MFN2 deficiency is actually associated with contributing to the UPR response as well as cellular apoptosis because MFN2 plays a role in suppressing PERK [80,81]. Previous studies have also demonstrated that ER-mitochondrial tethering can contribute to the upregulation of Drp-1 mediated fission via formation of constriction sites [82,83]. Recently, it was found that Drp1 is activated in experimental models for multiple sclerosis, and inhibition of its pathological hyperactivation is neuroprotective [84]. For the first time, we show that GA reduces mitochondrial fission activity and increases fusion activity in EAE mice, again strongly suggesting that GA targets ER stress and downstream mitochondrial mediators.

To further explore the effects of EAE and treatment of EAE with immunomodulating therapy on metabolism we examined expression of nicotinamide phosphoribosyltransferase (NAMPT), the rate limiting step in the NAD+ salvage pathway which has neuroprotective effects [53] and Sirt 1 and 3, members of the Sirtuin family which are NAD dependent protein deacetylases which are key metabolic sensors in the stress response [54]. Increasing SIRT1 activity, either by treatment with the Sirt activator, resveratrol [85] or by genetic overexpression [34], reduces the clinical and pathological severity of EAE. SIRT3 activates PGC-1α which stimulates mitochondrial biogenesis and is associated with ROS suppression and neuroprotection[86]. Sirt1 is of further interest because levels are reduced in peripheral blood mononuclear cells during relapses of MS[87] and were restored by treatment with GA[88]. Sirt3 changes are also implicated in MS in that levels were reduced in non-lesioned grey matter from MS brains[89]. We found that expression of NAMPT and Sirt 1 and 3 were reduced in EAE and restored or partially restored to normal levels by treatment with immunomodulating therapy with glatiramer acetate. This suggests that such therapy can restore cells to a more normal metabolic state.

To determine whether the reductions in ER stress, mitochondrial dysfunction and metabolic abnormalities induced by immunomodulatory therapy were associated with reductions in cellular death we examined apoptosis and changes in related pathways. The increased fission processes that we found in the EAE model can contribute to cellular apoptosis by the opening of BAX lined pores and release of cytochrome C [90], and our findings supported this. Using TUNEL staining we found that immunomodulating therapy with GA reduced the high levels of apoptosis is found in untreated EAE. We studied the expression of FOXO transcription factors implicated in the regulation of apoptosis [55] and found increases in EAE which were reversed by GA treatment. We examined the expression of Bax, a BCL-2 family member that promotes apoptosis by contributing to mitochondrial membrane pore formation[56] and found increased levels in EAE which were reversed by GA treatment. These results show that while EAE is associated with an increase in cell death, treatment with the immunomodulating therapy, GA, is associated with a reduction. Our results strengthen the idea that the mitochondrial and ER changes in EAE are part of a coordinated response to metabolic stress caused by inflammation and that immunomodulating therapy can reduce that stress.

We found mitochondrial dysfunction, endoplasmic reticulum stress and disrupted NAD metabolism in the spinal cords of EAE mice. For the first time, we show that GA can potentially reverse these pathological changes, and Figure 7 shows our proposed mechanism. Given that the direct effects of GA are thought to be in the peripheral immune system, it seems likely that the observed changes are an indirect immunomodulatory effect resulting in reduced inflammation in the spinal cord, and thus neuroprotective. The reduction in neuronal apoptosis in EAE upon GA treatment could be a result of ameliorated ER stress, improved mitochondrial function, and regulated NAD metabolism.

**Fig 7: Proposed neuroprotective mechanism underlying GA treatment in EAE**. During EAE, astroglia and microglia are activated by the infiltrated cells, which includes Th1 cells (T-cells, activated macrophages, B-cells, and Neutrophils) and contribute to a neuroinflammatory environment that damages neurons. As a result, this causes mitochondrial dysfunction and ER stress in neuronal cells. GA induces a shift towards the Th2 response in the periphery, and these Th2 cells cross the blood brain barrier into the CNS. This reduces the inflammatory environment and controls the synergistic ER stress response and mitochondrial dysfunction. By doing so, apoptotic activity in the CNS is downregulated, likely in the myelin producing oligodendrocytes. In turn, disease progression is slowed. CNS AG: CNS antigen; MHC: major histocompatibility complex; TCR: T cell receptor.

## Acknowledgements

This work was supported by grants from Teva Neuroscience, Inc. and the Department of Veteran’s Affairs

## Author contributions

C Bever and T Makar designed the study. T Makar induced the EAE in mice, and injected the mice with glatiramer acetate. T Makar and V Nimmagadda euthanized the mice and collected the tissue. V Nimmagadda, P Guda, and S Andhavarapu performed the immunohistochemical staining. P Guda did the western blot analysis. S Ray prepared tissue for electron microscopy, took the pictures, and provided the analysis. T Makar, S Andhavarapu, V Nimmagadda, V Gerzanich, and C Bever analyzed and interpreted the data. S Andhavarapu, T Makar, V Gerzanich, and C Bever drafted the manuscript. All authors critically reviewed and approved the final version of the manuscript.

## Abbreviations

MS: multiple sclerosis
CNS: central nervous system
EAE: experimental autoimmune encephalomyelitis
UPR: unfolded protein response
MRI: magnetic resonance imaging
ROS: reactive oxygen species
ER: endoplasmic reticulum
MAM: mitochondria associated membrane
NAD: nicotinamide dinucleotide
GA: glatiramer acetate
BBB: blood brain barrier
Ca2+: calcium ions
MOG: myelin oligodendrocyte glycoprotein
CFA: complete Freund’s adjuvant
PBS: phosphate-buffered saline
IL-17: interleukin 17
MBP: myelin basic protein
MFN-2: mitofusin-2
Fis-1: fission-1
DNM1-L: dynamin-like protein
PGC1-a: Peroxisome proliferator-activated receptor gamma coactivator 1-alpha
FOXO-1: Forkhead box protein 1
TFAM: mitochondrial transcription factor A
SIRT: sirtuin
NAMPT: Nicotinamide phosphoribosyltransferase
BAX: Bcl-2-associated X protein
CHOP: CCAAT-enhancer-binding protein homologous protein
PERK: protein kinase RNA-like endoplasmic reticulum
PINK1: PTEN-induced kinase 1
VDAC-1: Voltage-dependent anion channel 1
PACS2: Phosphofurin Acidic Cluster Sorting Protein 2
eIF2α: eukaryotic translation initiation factor 2 subunit alpha

